# Human sperm chromatin forms spatially restricted nucleosome domains consistent with programmed nucleosome positioning

**DOI:** 10.1101/481028

**Authors:** Mei-Zi Zhang, Xiao-Min Cao, Feng-Qin Xu, Xiao-Wei Liang, Long-Long Fu, Fang-Zhen Sun, Xiu-Ying Huang, Wei-Hong Huang

## Abstract

In human sperm, a fraction of its chromatin retains nucleosomes that are positioned on specific sequences containing genes and regulatory units essential for embryonic development. This nucleosome positioning (NP) feature provides an inherited epigenetic mark for sperm. However, it is not known whether there is a structural constraint for these nucleosomes and, if so, how they are localized in a three-dimensional (3D) context of the sperm nucleus. In this study, we examine the 3D organization of sperm chromatin and specifically determine its 3D localization of nucleosomes using structured illumination microscopy. A fraction of the sperm chromatin form nucleosome domains (NDs), visible as microscopic puncta ranging from 40 μm to 700 μm in diameter, and these NDs are precisely localized in the postacrosome region (PAR), outside the sperm’s core chromatin. Further, NDs exist mainly in sperm from fertile men in a pilot survey with a small sample size. Together, this study uncovers a new spatially restricted sub-nuclear structure containing NDs that are consistent with NPs of the sperm, which might represent a novel mark for healthy sperm in human.

## Introduction

In animal sperm, genomic DNA is mainly packaged by sperm nuclear basic proteins (SNBPs), which includes protamines, protamine-like proteins, and H1 histone-type proteins (1,2). In mammalian sperm chromatin contains somatic histones to form nucleosomes and SNBPs to form nucleoprotamine. The retained nucleosomes package sequence-specific DNA fragments (3,4); whereas nucleoprotamine-containing chromatin forms the core chromatin. It is known that epsilon and gamma globin genes, which are expressed in the sperm, are embedded in nucleosomes in the sperm, but beta- and delta-globin genes, which are silent, are not associated with nucleosomes (4).

In mature human sperm, most of the retained nucleosomes are enriched in certain loci of the genome, such as imprinted genes and HOX genes, which are crucial for embryo development and can serve as epigenetic mark (5). This nucleosome-associated epigenetic information, such as histone H3 lys4 demethylation, in human and mouse sperm has been shown to be involved in spermatogenesis and cellular homeostasis, and has been used to mark developmental regulators by histone H3 Lys27 trimethylation (6).

DNA sequences embedded in retained nucleosomes in human sperm have been mapped, which are closely related to the established DNA-methylation-free zone in the genomes of early embryos. From an evolutional point of view, the selective pressure on certain base compositions helps to retain specific nucleosomes, allowing successful transmission of specific paternal epigenetic information to the zygote. (7, 8). While in mouse sperm only 1% genome is embedded into nucleosomes, and in human sperm the nucleosome-associated chromatin constitutes almost 15% of the human genome retaining specific nucleosomes in the sperm as an epigenetic feature conserved over generations is consistent for both human and mouse (9, 10).

Using proteomic mapping of human sperm, it has been found that retained nucleosomes constitute additional layers of epigenetic information (11). Recent studies have shown that nucleosomes in human sperm are specifically positioned at transcription start sites (9,11). In genome-wide surveys of somatic cells and sperms, nucleosomes are preferentially positioned at internal exons; however, this is independent of their epigenetic modification status such as histone methylation and acetylation. This type of chromatin structure may involve general transcriptional regulation, exon recognition and splice site selection (12). However, certain conclusion about genome organization, based solely on computational analyses, has been challenged (13). New experiments to determine, or validate, structural features of the genome are needed.

While numerous studies suggest that nucleosome positioning in human sperm is important for early embryonic development and paternal epigenetic inheritance (5-8), specific nucleosome positioning has yet to be experimentally confirmed. Further, the information regarding any 3D localization of nucleosome domains is entirely unknown. In this study, we determine the 3D positioning of nucleosomes in human sperm by using super-resolution structured illumination microscopy (SIM). We conclude that in human sperm, histone-containing chromatin forms nucleosome domains (NDs), which are specifically localized at PAR and represent a structural feature of chromatin organization in the sperm. Further, NDs were mainly found in the sperm from sterile men in a pilot survey, indicating that NDs might be associated with healthy status of sperm in human.

## Result

We first obtained healthy sperm samples for this study, the quality of sperms used in the study is shown in table I. Based on WHO morphology guidelines, the fresh semen sample contains a high percentage (68%) of sperms exhibiting normal morphology (14, 15), indicative of fertility. After gradient centrifugation through Puresperm 100 media, the purified sperm fraction has a 100% motility rate, a 99% progression and a 100% normal morphology.

**Table I.** comparsion of sperm parameters after density-gradient centrifugation method(puresperm100)for the fertile man

|  | <u>control (fresh semen)</u> | <u>PureSperm</u> |
| --- | --- | --- |
| Concentration ($\times 10^6$)/ml | 45 | 21 |
| Total motility (%) | 42 | 95 |
| Normal morphology (%) | 68 | 96 |
| Progressive motility (%) | 37 | 98 |

We embedded purified sperms in a 20% gelatin solution, and then in optimal cutting temperature compound (OCT compound), which maintains the 3D structure of the sperm, allowing 3D structure to be determined by SIM microscope. Cryo sections of the OCT-embedded sperm block were obtained at the thickness of 4 μm, which were subjected to immunostaining using histone H4 antibody alone or in conjunction with protamine 1 antibody. The slides were examined by SIM and images were analyzed by softWoRx Suite software (GE, USA). Histone H4-positive signals indicate the presence of nucleosomes. The staining patterns can be classified into two major types with type I containing sparse puncta of H4 signals and type II containing aggregated puncta of H4 signals (Figure 1A and 1B, Video S1 and S2). These distinct puncta indicate that nucleosomes in human sperm form specific structures, such as NDs, because the size of these puncta are much larger than that of a single nucleosome (around 10 nanometer). Type I structure represents the dominant type, whereas type II is occasionally seen. These types of histone H4-positive NDs are found in most of the sperms, which are also found by suing histone H2B antibody (Figure S1). The statistical results are shown in Table II.

**Table II.**
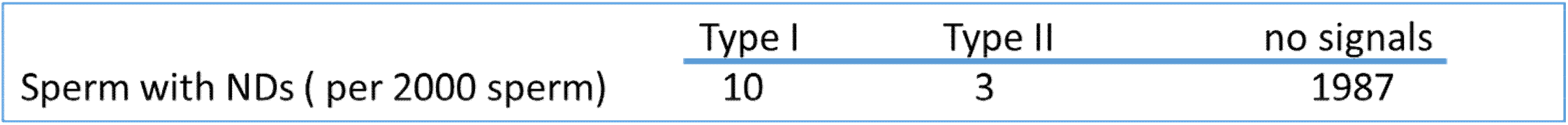
the clasification of NDs

|  | Type I | Type II | no signals |
| --- | --- | --- | --- |
| Sperm with NDs ( per 2000 sperm) | 10 | 3 | 1987 |

**Figure 1.**
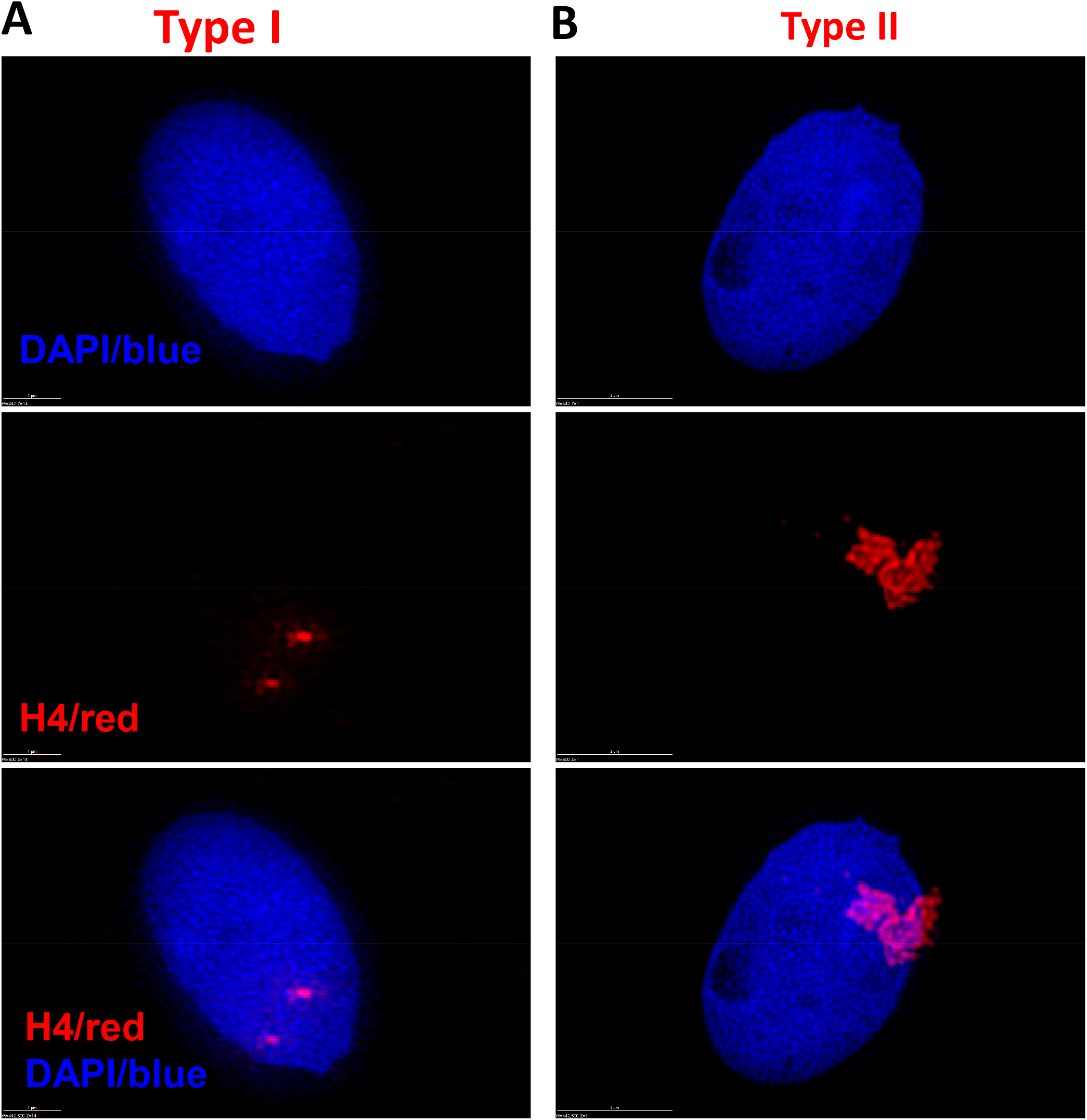
Nucleosomes in human sperm localize in PAR with sparse or aggregated dots to form NDs. (A) Type I NDs with sparse dots. The diameter of NDs ranges from 40-700 μm. (B) Type II NDs with aggregated dots. Type I is popular, while type II is minor.

Interestingly, NDs are strictly localized around the PAR and are outside of zone of core chromatin (Figure 2 and S2, Video S3 and S4). The size of NDs arranges from 40 to 700 μm in diameter. The number of NDs in a sperm ranges from 9-67, but the major NDs only range from 3-19 (Table III).

**Table III.** The parameters of NDs

| sample number | size (nanometer, nm) | numbers | major NDs |
| --- | --- | --- | --- |
| 1 | 80-700 | 30 | 16 |
| 2 | 60-630 | 33 | 15 |
| 3 | 50-400 | 34 | 14 |
| 4 | 60-600 | 39 | 7 |
| 5 | 60-500 | 31 | 7 |
| 6 | 80-630 | 31 | 7 |
| 7 | 60-500 | 49 | 19 |
| 8 | 43-500 | 11 | 3 |
| 9 | 80-500 | 15 | 5 |
| 10 | 40-400 | 67 | 10 |
| 11 | 60-300 | 20 | 6 |
| 12 | 40-400 | 40 | 9 |
| 13 | 40-500 | 42 | 9 |
| 14 | 40-400 | 30 | 15 |
| 15 | 90-300 | 9 | 4 |

**Figure 2.**
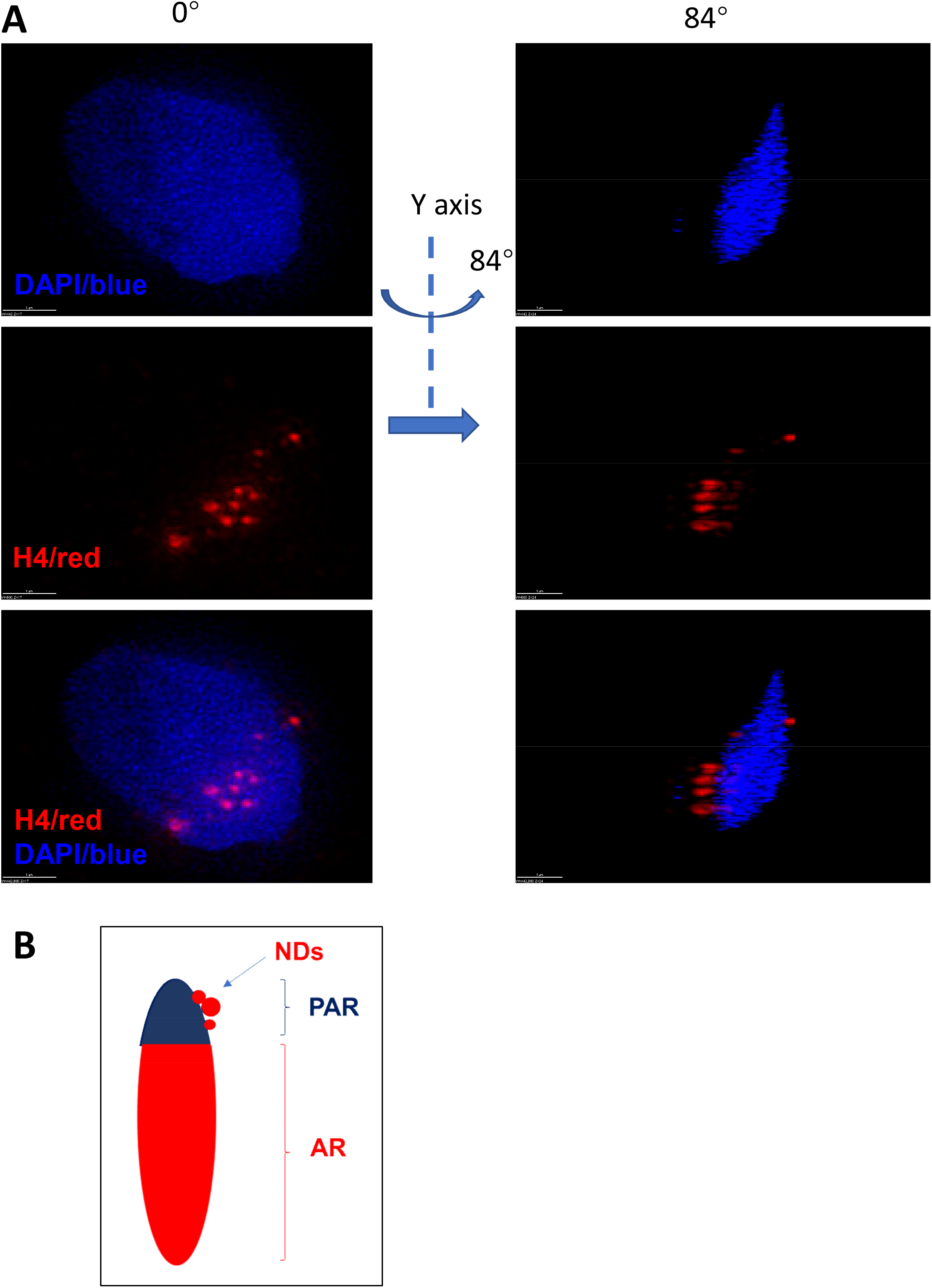
By use of SIM the 3D positioning of NDs was shown. (A) After rotation around Y axis, NDs was shown to be localize outside of core chromatin in PAR. The NDs were not shown to be inside of sperm chromatin. (B) A diagram was shown. NDs are positioning outside of major dense sperm chromatin.

To further localize the NDs in sperm chromatin, we examined those sections spanning the sperm inner chromatin, and determined whether NDs are also localized in the inner sperm chromatin. Protamine 1 stains the sperm core chromatin. On longitudinal sections cut through the acrosome region (AR) or PAR, we found that those sections spanning the AR are protamine 1-positive and H4-negative. However, those sections spanning the PAR are H4-positive only outside the core sperm chromatin. H4 positive signals are never found in the inner chromatin which is stained positive with protamine 1 (Figure 3, 4, and S3, Video S5, S6 and S7).

**Figure 3.**
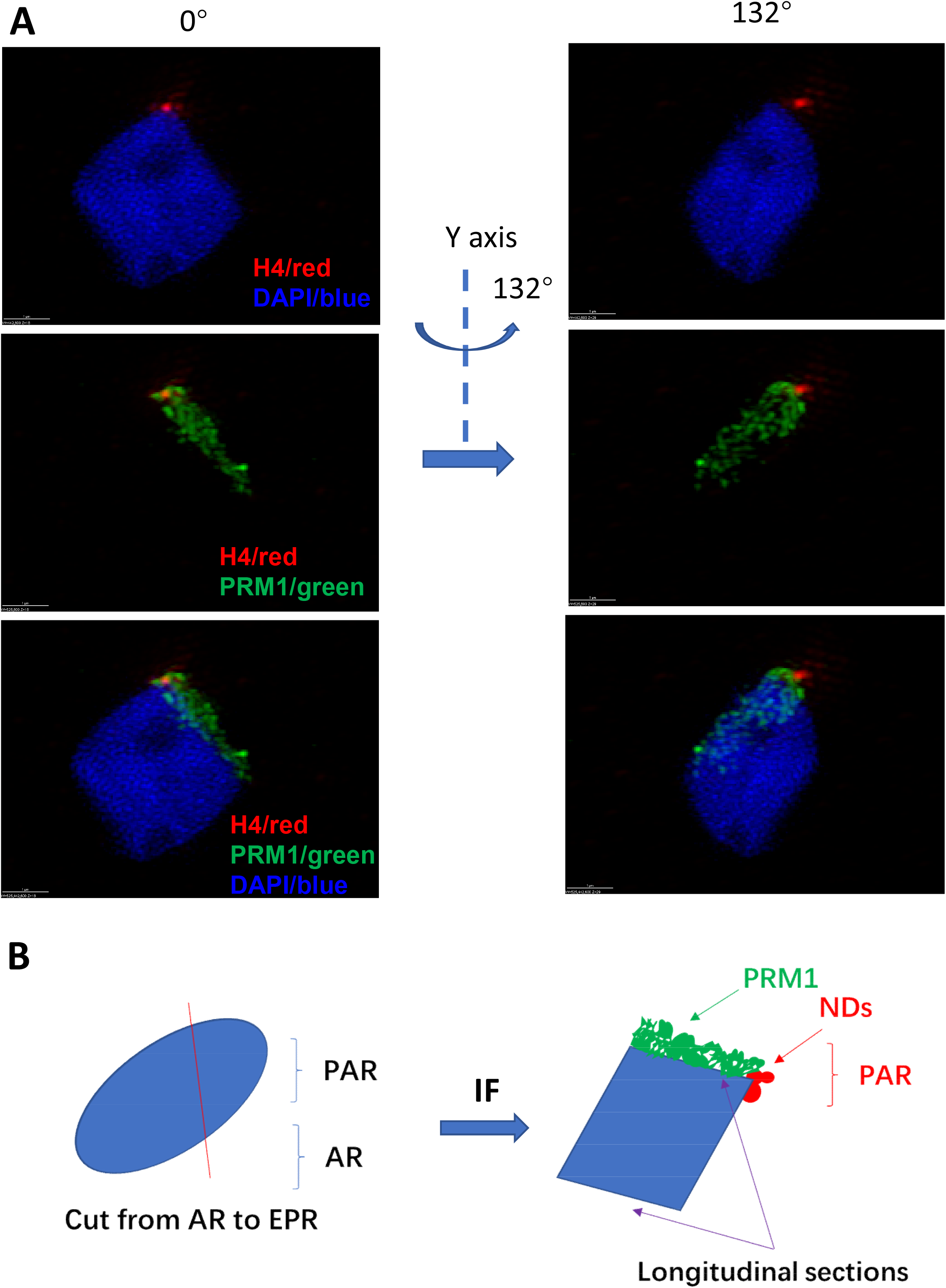
After cutting through AR and PAR, localization of NDs was shown. (A) After this sperm was cut on both sides, the inner side stick to the slide without staining by antibodies but the outside side was stained by both H4 and protamine 1 antibodies. H4 signals were confined in the PAR and outside of inner chromatin, but not in inner chromatin. However, protamine 1 signals ranged from the AR to PAR in inner chromatin in longitude section. (B) A diagram was shown. NDs are not localized in the inner chromatin but outside of sperm chromatin in the PAR.

**Figure 4.**
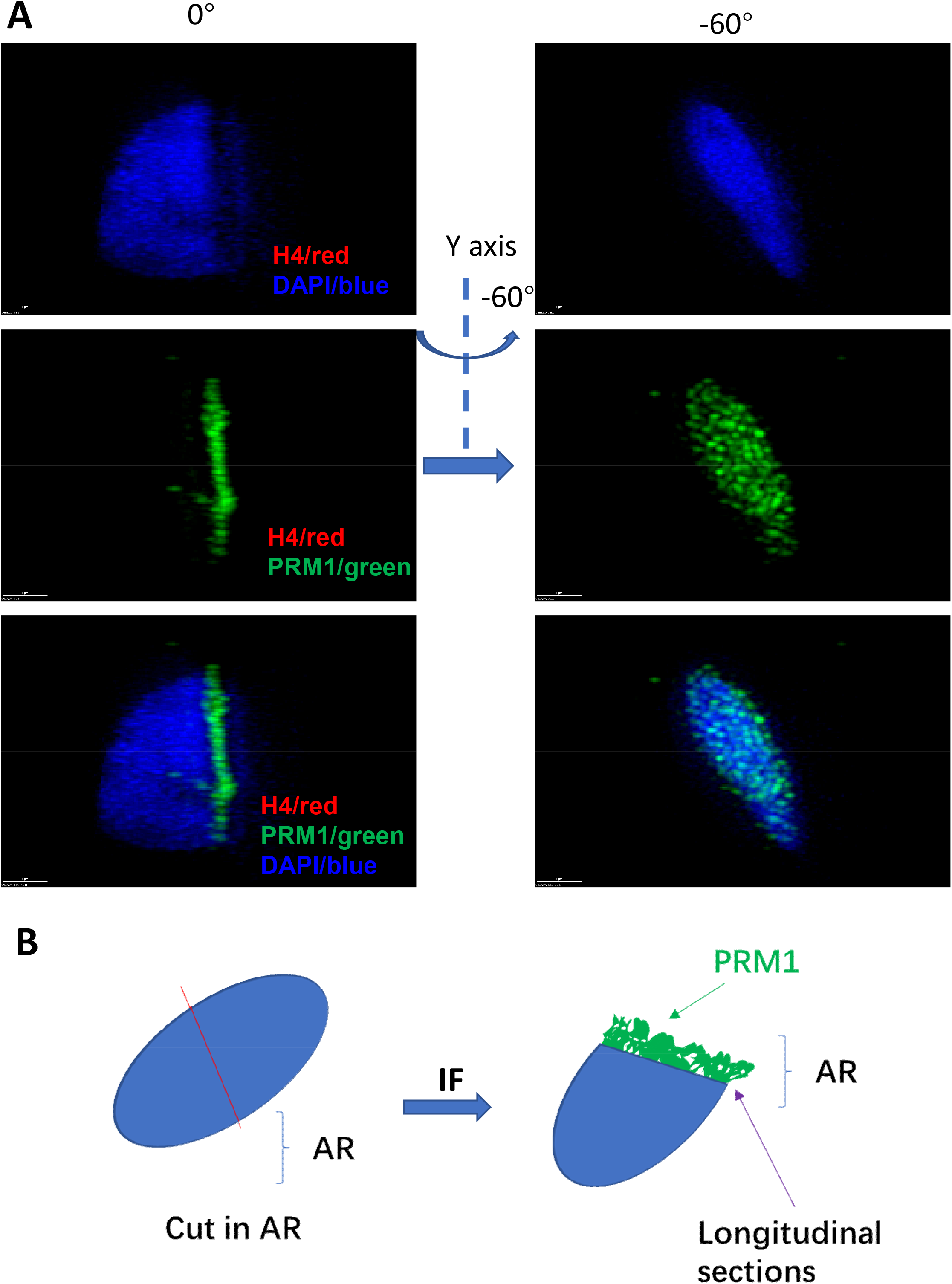
After cutting through AR NDs was not detected. (A) This sperm was cut in AR. After staining by both H4 and protamine 1 antibodies, H4 signals were not detected, while protamine 1 signal was full of the inner chromatin in longitude section. (B) A diagram was shown. NDs were not detected in inner and outside of sperm chromatin in AR.

These data show that nucleosomes in human sperm form NDs which are specifically localized, proximal to the PAR. This is consistent with studies suggesting that nucleosomes in human sperm form on certain specific sequences of the genome (3, 5, 16). It has been proposed that specific nucleosome positioning (NP) in human sperm genome is important in guiding programmed gene expression in early embryonic development and may represent a form of epigenetic inherence (5,7). We thus propose that NDs observed in our study here may reflect NP in the sperm. Further, we collected six infertile samples and six fertile samples, and surveyed the distribution of NDs in these samples. Interestingly, none of the six infertile samples exhibit any form of NDs, but all the six fertile samples contain sperm with NDs (table IV). The results imply that NDs in human sperm might be relative the healthy status of sperm.

**Table IV.** semen quality of the sperm donars in fertitle and sterile groups

|  | fertile group |  |  |  |  |  | sterile group |  |  |  |  |  |
| --- | --- | --- | --- | --- | --- | --- | --- | --- | --- | --- | --- | --- |
|  | Patient 1<br>(32 yr) | Patient 2<br>(34 yr) | Patient 3<br>(31 yr) | Patient 4<br>(29 yr) | Patient 5<br>(34 yr) | Patient 6<br>(36 yr) | Patient 1<br>(36 yr) | Patient 2<br>(35 yr) | Patient 3<br>(39 yr) | Patient 4<br>(30 yr) | Patient 5<br>(33 yr) | Patient 6<br>(34 yr) |
| Semen volume (ml) | 3.6 | 3.9 | 4.2 | 3.3 | 4.3 | 3.1 | 3.1 | 2.9 | 3.4 | 3.8 | 5.6 | 4.9 |
| Total motility (%) | 46.58 | 59.23 | 56.58 | 42.36 | 52.84 | 57.24 | 49.43 | 61.37 | 52.69 | 63.28 | 60.63 | 48.34 |
| Sperm concentration ( x 10<sup>6</sup> per ml) | 48.98 | 84.41 | 63.72 | 58.74 | 48.35 | 51.53 | 43.56 | 57.94 | 67.28 | 61.63 | 49.68 | 54.72 |
| Progressive motility (%) | 43.36 | 50.11 | 40.61 | 39.57 | 44.75 | 54.98 | 39.45 | 45.37 | 48.94 | 36.08 | 34.78 | 53.09 |
| Morphologically normal sperm (%) | 31.25 | 39.76 | 40.69 | 48.58 | 50.87 | 56.78 | 42.46 | 31.52 | 48.12 | 44.36 | 47.07 | 35.18 |
| Pregnant | yes | yes | yes | yes | yes | yes | not | not | not | not | not | not |
| Sperm with NDs ( per 2000 sperm) | 8 | 9 | 13 | 11 | 10 | 7 | 0 | 0 | 0 | 0 | 0 | 0 |
| NP | PAR | PAR | PAR | PAR | PAR | PAR | not | not | not | not | not | not |

Taken together, this study indicates that NDs, which is specifically localized and is proximal to the PAR, could represent a new type of 3D structural feature in human sperm. Further, in a pilot survey the presence of NDs is shown to be mainly in fertile sperm, indicating that NDs might be relative to the healthy status of sperm in human.

## Discussion

Though where nucleosomes are localized in mammalian sperm genome remains controversy, it is a generally consistent view that retained nucleosomes in the sperm influence embryonic development and epigenetic inheritance (3, 5, 17, 18).

It is hypothesized that sequence-specific NP facilitates sperm chromatin decondensation (SCD) upon fertilization, and that the position represents the start site of SCD (3). SCD occurs immediately after membrane fusion of gametes (19) and the site in the sperm for membrane fusion is also the start site of SCD, which has been shown to be around the PAR (20, 21). In this study, we present evidence for NDs in the sperm, and show that NDs form around the PAR.

Previous studies have shown that nucleosomes in human sperm package specific DNA sequences, which provide epigenetic marks and are required for embryonic development (3, 5, 6). In this study, our data demonstrate that in human sperm, nucleosomes form distinct NDs which are specifically localized to where NP is suggested to exist. This would indicate that both NDs and NP in the sperm are the results of important,programmed events during spermatogenesis, and support embryonic development after fertilization.

Strikingly, NDs are not found in inner chromatin. As protamines in core chromatin might handle the binding of histone antibodies and previous studies had showed that without SCD immunostaining for protamines and histones in human and mouse sperm was failed (22, 23), single nucleosomes might still sparsely distribute in inner chromatin. However, as NDs are large units and unlikely are fully inhibited by protamines during immunostaining, it is very likely that NDs are not localized in inner chromatin.

As nucleosomes of mammalian sperm will transmit into zygote (22), paternal epigenetic information on NP should be programmed and transferred to their children. In previous study (16), when MNase was used to digest human sperm chromatin, dinucleosomal bands and nucleosome ladders were found. The researchers hypothesed that nucleosome retention occurs in blocks with adjacent nucleosomes (16). Our data is highly consistent with those observations as NDs are programmed to precisely aggregate in the PAR and outside of inner chromatin.

Infertility affects around 15% couples around the world, and male infertility accounts for 40-50% of it (24, 25, 26, 27). In this study our data indicates that NDs and NP might constitute additional layers of epigenetic information in human sperm, which then could be a new target in diagnosing male infertility.

## Materials and Methods

### Specimen collection

All procedures were in accordance with institutional guidelines on Human Subjects in Experimentation of the Institute of Genetics and Developmental Biology (CAS, China), Tianjin First Central Hospital, and National Research Institute for Family Planning. Semen samples were collected by manual masturbation from consenting men with known fertility (having a child within the past two years) or with known sterility (without a child in history by natural conception, but the semen parameters are normal. Their partner had not conceived in IUI, IVF and ICSI using their semen, but had conceived in IUI by use of healthy donor’s semen.).

### Specimen processing

After liquefaction at 37°C for 60 minutes in an air incubator with 5% CO2, seminal plasma was removed by centrifugation at 300g for 10 minutes, through Enhance S-Plus Cell Isolation Media (mHTF, Vitrolife, https://www.vitrolife.com). The cells were washed in Modified Human Tubal Fluid medium (Irvine Scientific, www.irvinesci.com) once, and pelleted by centrifugation at *700g* for 10 minutes. The sperm were resuspended in mHTF at a high concentration (about 200 × 10^6^ per ml) fixed by 4% paraformaldehyde (pH 7.4) in 1 x PBS for overnight at 4°C. After removing excess paraformaldehyde with PBS washing, the fixed sperm was subjected for gradient sucrose infiltration. Further, after removal of excess sucrose from the fixed sperm, it was embedded in OCT compound. After the fixed sperm are frozen, it was placed into plastic bags, sealed and stored at −20°C.

### PureSperm density gradient centrifugation

By using a PureSperm 100 discontinuous density gradient (Nidacon, Gothenburg, Sweden), a two-layer gradient was prepared by dilution solutions of 40% and 80% PureSperm. After the whole media was pre-warmed to 37°C, liquefied semen sample was placed on top of the upper layer and centrifuged at 300g for 20 minutes following the product’s instructions. The pellet was washed twice by wash solutions and recovered; aliquots were used for storage at liquid nitrogen or used for a routine semen analysis according to the World Health Organization guidelines (2010, 5th).

### Cryopreservation of spermatozoa and thawing

After purified sperm was diluted in Sperm Maintenance Medium (SMM, Irvine Scientific, USA) at the ratio of 3:1, the mixture was transferred to 0.5 ml straw (PETG sperm 0.5 ml-clear straw, Irvine Scientific, CA, USA). That was suspended for 40 minutes in liquid nitrogen vapor, and then plunged into liquid nitrogen for long storage.

After the straw was pulled out from liquid nitrogen, it was immersed immediately into prewarmed mineral oil at 38°C. After warming, sealer of the straw was removed, and sperm was washed once by Sperm Washing Medium (Irvine Scientific, CA, USA).

### Immunohistochemistry

After thawing, sperm was fixed in 4% paraformaldehyde in phosphate buffered saline (1 x PBS; pH 7.4) for 30 minutes at room temperature or overnight at 4°C. The fixed sperm was centrifuged at 3,000g for 5 minutes, and the pellet was embedded into 20% gelatin in water at 42°C. After the samples were solidified at 4°C for 1hour, they were embedded in OCT compound (Fisher Scientific, Pittsburgh, PA) and frozen in liquid nitrogen. Frozen blocks of sperm were then cytosectioned at 4 μm thickness (Leica CM3050 S, Leica, Germany),and the slices were put on slides and dried at room temperature for at least 2 hours. The slides were further washed by PBS with 0.1% tween-20 and 0.01%Tx-100, they were permeabilized with 0.5% Triton X-100 in PBS at room temperature for 1 hour. After washed by PBS with 1% bovine serum albumin (BSA) twice, they were blocked in 1% BSA - supplemented PBS in for 1 hour at RT. After incubated overnight at 4°C in the appropriate antibodies (Histone-H4 Rabbit Polyclonal antibody (1: 100, Proteintech Group, 16047-1-AP), Histone-H2B Rabbit Polyclonal antibody (1: 100, Proteintech Group, 15857-1-AP) or Protamine 1 Antibody (A-17) (1:200, Santa Cruz, sc-23107)) diluted in 1% BSA-supplemented PBS, they were washed in PBS containing 0.1% Tween 20 and 0.01% Triton X-100 for 5 minutes each three times. Then, a secondary antibody labeled with Alexa Fluor 594 or 488 (Goat Anti-Rabbit IgG-Alexa Fluor 594, ab150080; Goat Anti-Rabbit IgG-Alexa Fluor 488, ab150077; Donkey Anti-Goat IgG-Alexa Fluor 488, ab150129) was used as the fluorescent reporter for the primary antibodies. Then they were post-fixed with 4 % formaldehyde in PBS for 30 min at RT. After washed for 5 min by PBS with 0.1% tween-20 and 0.01% Tx-100 three times, they were stained by DAPI (1 μg/ml) for 20 minutes at RT. After washed by PBS containing 0.1% Tween 20 and 0.01% Triton X-100 for 5 min each time twice, they were drained and mounted in ProLong Gold Antifade Reagent (Life Tech). The sealed slides were used for 3D study by SIM.

### 3D-SIM super-resolution microscopy and image analysis

SIM was performed on the DeltaVision OMX V3 imaging system (Applied Precision) with Plan Apochromat 100x/1.46 Oil objective, 1.6x lens in the detection light path and Andor iXon 885 EMCCD camera; Z-stacks were acquired with 125 nm intervals; In three-color images, channels (405, 488, and 593 nm) were acquired sequentially using standard single-band filter sets. SIM image stacks and raw SIM images were analyzed by softWoRx 5.0 (Applied Precision). Further, reconstructed images were rendered in three dimensions also by softWoRx 5.0, and linear adjustments to brightness were performed on 3D reconstructions for better contrast.

### Quantification of nucleosome domains and positions

The nucleosomes domains and positions were analyzed by softWoRx 5.0 software. Measurement module was used to calculate the size of ND. In briefly, using “measure tool”, the method is “multiple segment” and the units is “micrometers”. Manually selecting a start point and an end point on one ND, the software will return the length between the two points. This length was then considered to be the diameter of this ND.

## Acknowledgement

We are very grateful to professor Li-Na Wei (Department of Pharmacology, Medical School, University of Minnesota, MN, USA) on revising the whole manuscript. We also are grateful to Shuo-Guo Li for access to the DeltaVision OMX super-resolution microscope (SIM) at the Institute of Biophysics. This work was supported by a grant from Ministry of Science and Technology (2012CB944903).

---

**Figure S1.**
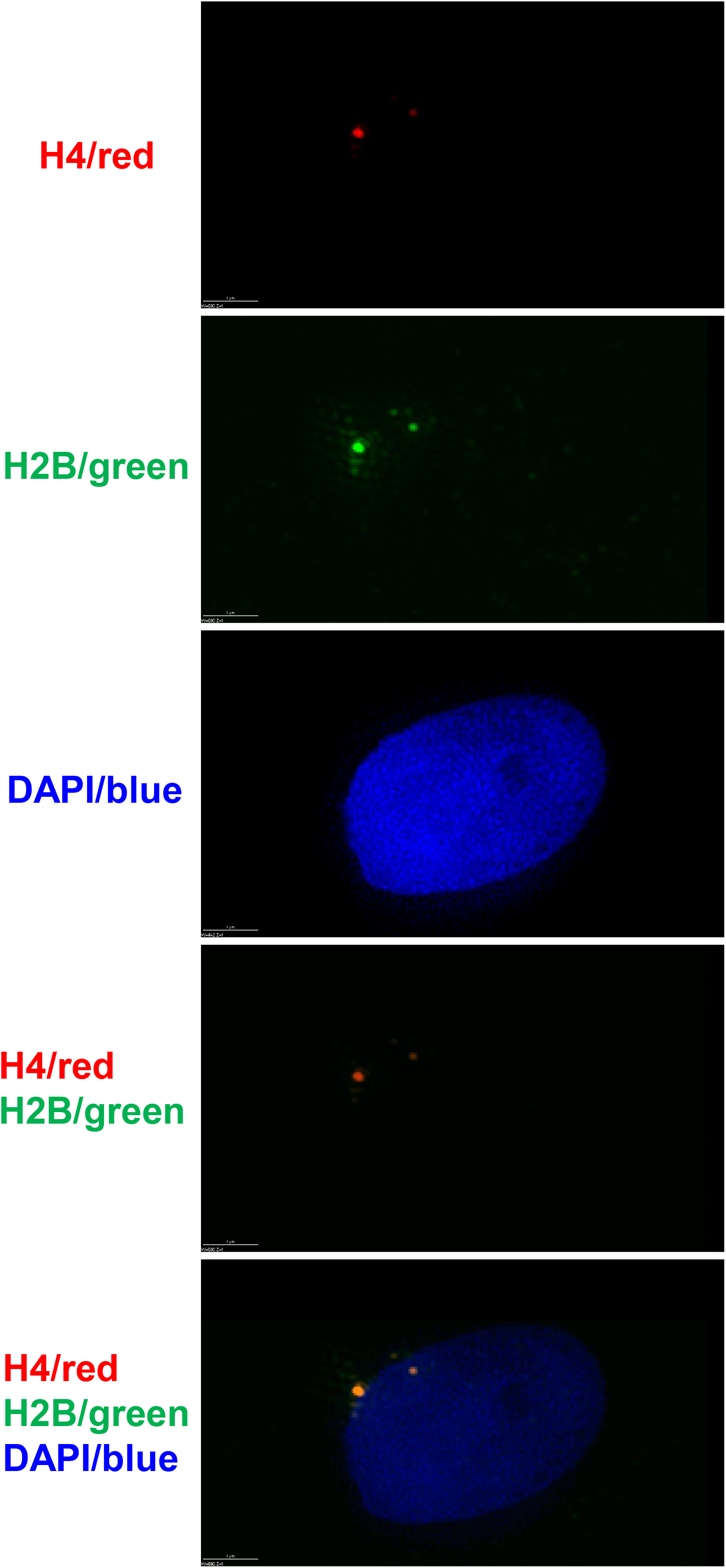
NDs were also detected by use of histone H2B antibody. Similar results were obtained by use of histone H2B antibody. NDs with sparse dots were shown outside of sperm chromatin in PAR.

**Figure S2.**
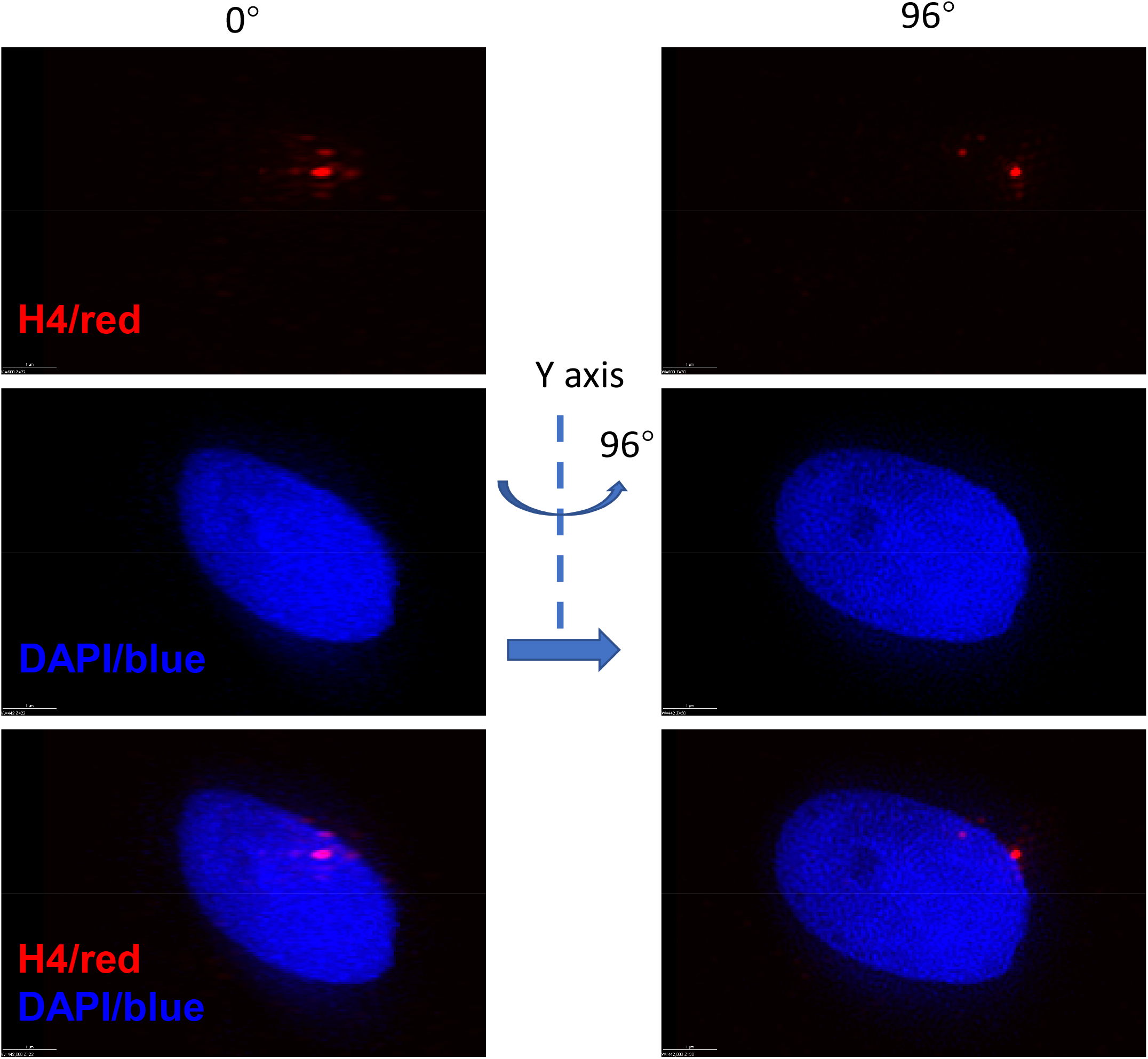
NDs was shown to be exactly localize outside of core chromatin in PAR. Similar results were obtained in another sample. NDs with sparse dots were shown outside of inner chromatin in PAR. This data is similar to Figure 2.

**Figure S3.**
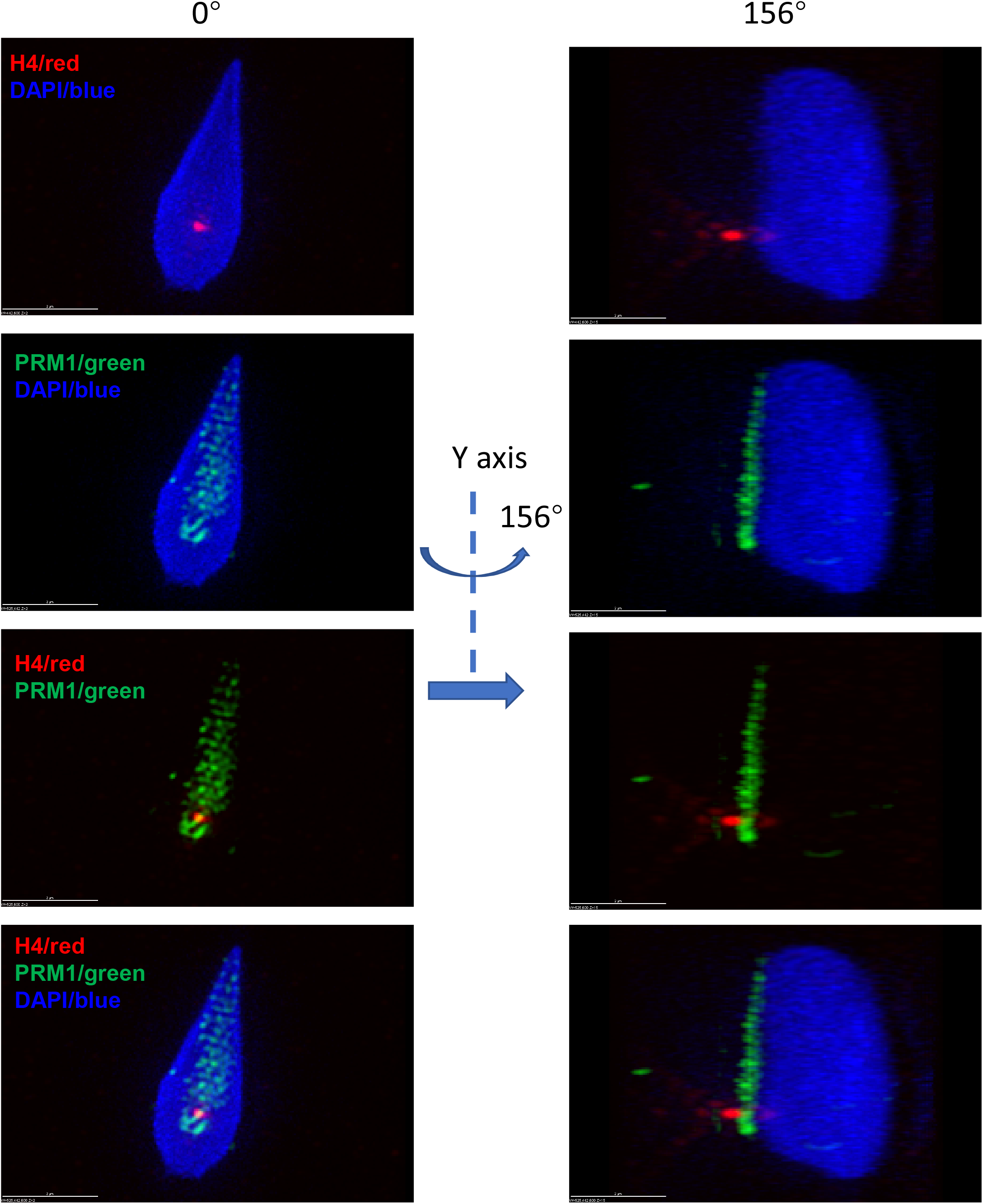
NDs was confined in the PAR and outside of inner chromatin, but not in inner chromatin. Similar results were obtained in another sample to show that NDs localized outside of sperm chromatin but not in the inner sperm chromatin. NDs with sparse dots were shown outside of inner chromatin in PAR, but not in inner chromatin. This data is similar to Figure 3.

